# Spectral detection of condition-specific biological pathways in single-cell gene expression data

**DOI:** 10.1101/2023.03.12.532317

**Authors:** Wee Loong Chin, Leonardo Portes dos Santos, Michael Small, W. Joost Lesterhuis, Timo Lassmann

## Abstract

Single cell RNA sequencing is an ubiquitous method for studying changes in cellular states within and across conditions. Differential expression (DE) analysis may miss subtle differences, especially where transcriptional variability is not unique to a specific condition, but shared across multiple conditions or phenotypes. Here, we present CDR-g (Concatenate-Decompose-Rotate genomics), a fast and scalable strategy based on spectral factorisation of gene coexpression matrices. CDR-g detects subtle changes in gene coexpression across a continuum of biological states in multi-condition single cell data. CDR-g collates these changes and builds a detailed profile of differential cell states. Applying CDR-g, we show that it identifies biological pathways not detected using conventional DE analysis and delineates novel, condition-specific subpopulations in single-cell datasets.

## 1 INTRODUCTION

Single cell sequencing allows gene expression profiles to be mapped to individual cells at scale. It has enabled the discovery of new therapeutic targets^1,2^ and improved our understanding of developmental processes.^3,4^ Reinforcing the utility of this technology, many single cell atlases have become available in the public domain^5,6^ For example, the Human Cell Atlas project^6^ characterises cell identity across all human organ systems and the COVID atlas is a single-cell lung atlas of lung tissue from patients with lethal COVID-19.^7^ Efforts have also been made to construct atlases combining single cell datasets across multiple studies – for instance, the immune cell atlas contains more than 500,000 cells from 217 patients and 13 cancer types.^8^ Transcriptomic profiling of cancers such as metastatic lung adenocarcinoma has afforded a deep understanding of tumour cell state.^9^ Integrative analysis of data from these large single cell atlases is a promising avenue to understand health and disease states.

A defining feature of these datasets is the presence of multiple samples, conditions and cell states therein. To analyse such data, DE is the most accessible first-pass analysis. However, there are several challenges to conventional DE analysis with such complex data. The number of comparisons grows with the number of conditions and populations analysed^10,11^ Additionally, DE detects statistically significant changes in average expression, so it does not capture more nuanced transcriptional heterogeneity such as differential co-expression^12^ and differential variation^13^ which may nevertheless be an important contributor to cellular identity. Multiple ways have been proposed to address this challenge posed from DE analysis when applied to complex, multi-condition datasets. For instance, phenotype information can be used to guide the selection of condition-specific genes. DIscBIO,^14^ a biomarker discovery pipeline, applies a decision-tree classifier on these refined lists to distinguish different conditions. Alternatively, genes which discriminate between conditions can be selected iteratively using wrapper methods, yielding a feature set of condition-specific genes^15^ Another strategy to interpret complex single cell datasets is to analyse cells in the entire dataset irrespective of condition, creating a representation of global transcriptional variation. Illustrating this strategy, SCENIC,^16^ an unsupervised, random-forest approach, uses gene expression information to construct a gene regulatory network across single cells and identifies gene sets associated with the activity of transcription factors, which can then be used to describe their condition-specific activation. In contrast, PAGODA2^17^ and Vision^18^ exploit low-dimensional feature spaces representing gene expression across conditions, which, downstream, allows transcriptional activity between conditions to be described in terms of differential signature activity of known gene sets.

Spectral techniques are another broad class of methods used to study single cell datasets and are widely applicable to multi-condition single cell data. Methods such as principal component analysis (PCA)^19^ and canonical correlation analysis (CCA)^20^ are used to pre-process and integrate single cell datasets. Matrix factorisation strategies such as consensus negative matrix factorisation (NMF)^21^ have also been used to study transcriptional variation in single cell data. These techniques decompose gene expression matrices into lower-dimensional spectra, where each spectra captures a component of variance which can be regarded as a gene expression program. By decomposing individual cell expression profiles into these programs, matrix factorization techniques allow shared and differential transcriptional effects to be captured simultaneously during analysis.^21^ Recently, in the field of complex network analysis, a novel mathematical framework was proposed to simultaneously characterize shared structures within and between a large number of networks.^22^ CDR (concatenate-decompose-rotate) is a fast and scalable algorithm supported by a transparent and straightforward mathematical theory rooted in Factor Analysis. We hypothesized that CDR, which leveraged spectral factorisation, could be well suited to detect both shared and differential gene expression programs between conditions in multicondition single-cell sequencing data. Applying CDR to single cell data, we applied singular value decomposition (SVD)^23^ directly to condition-specific gene co-expression matrices. Following this, we further transformed the resulting spectra from SVD decomposition using orthogonal-varimax^24^ to enhance the detection of condition-specific variation.

Here, we describe our extension of the CDR mathematical framework to multi-condition single cell data which we call CDR-g (CDR-genomics). To extend CDR, we implement a parameter-free gene selection strategy using a permutation filter,^25^ operating on varimax-optimised spectra. This permutation filter creates a dictionary of gene sets which describe both shared and differential expression programs between multiple conditions. The feature selection strategy allows us to rank individual genes in each gene set based on z-scores derived via permutation importance.^25^ Finally, we map the activity of these gene sets back to individual single cells using single sample gene set enrichment analysis (ssGSEA).^26^

## 2 MATERIALS AND METHODS

### 2.1 Overview of algorithm

For each condition *m* in a single cell dataset, a coexpression matrix is constructed for each condition using Pearson’s correlation coefficients. SVD is applied to the concatenated product of all *m* coexpression matrices.^22^ For *n* genes across *m* conditions with a choice of *N* factor loadings, this produces a factor loading matrix of dimension (mn) × *N.* Orthogonal varimax is applied to this loading matrix and the matrix retains its dimensions after this matrix transform. See^22^ for the mathematical theory supporting that step.

Computing the complete decomposition to produce the factor loading matrix is the most computationally intensive step of the CDR-g algorithm. To scale with single cell data, CDR-g uses truncated-SVD, which allows the user to specify the number of factors calculated during this decomposition. In CDR-g, we compute up to 2000 factor loadings or the number of factor loadings which captures 95% of variance across the input datasets.

For each rotated factor loading in the factor loading matrix, genes showing the highest differences in factor loadings scores show differential co-expression between conditions. We therefore designed a test statistic to extract these genes (features) based on a permutation filter.^25^ For each factor loading, we calculate the absolute difference in loading scores for each gene j across each of the m conditions. The rotated factor loading matrix is permuted and the test statistic recalculated. The number of times *nj* the test statistic of the random permutation is at least as large as the test statistic is used as a measure of feature strength. Strong features that warrant inclusion in a gene set have a *nj* ≤ 0.05*B*, where *B* is the number of permutations. For each gene, ranking is performed using z-scores, which measures the distance between the gene’s test statistic and the distribution of the test statistic of a random permutation.

CDR-g is written in python3, using established frameworks within the scientific computing stack (dask-ml, scipy, dask, pandas and numpy). The SVD algorithm used in the dask-ml library is a stochastic algorithm.^27^

### 2.2 CDR-g workflow and dataset pre-processing

A typical workflow with CDR-g comprises three steps. First, the CDR algorithm is used to identify condition-specific gene expression programs. Secondly, functional annotation of gene sets is performed. Finally, single sample gene set enrichment (ssGSEA) is used to map gene expression programs to individual cells, thus allowing identification of differential proportions and subpopulations of cells within each condition with testing of statistical significance. As input, CDR-g receives gene expression data in the form of UMI, TPM, or FKPM. The output of CDR-g is a set of ranked gene lists. Each gene list can be considered a gene expression program relevant to one or more conditions in the dataset.

We recommend the following steps before running CDR-g on a single-cell dataset. For gene filtering, we applied gene filtering thresholds based on those found in SCENIC.^16^ Briefly, two sequential filters are used: one based on the total number of counts of the gene and based on the minimal number of cells in which a gene s detected. We also scale and log-transform expression data as a variance-stabilising strategy prior to construction of gene coexpression matrices. When combining two datasets, highly variable genes were identified from each dataset using the “find variable genes” function in SCANPY^28^ and the set of intersections of these genes used for downstream analysis.

### 2.3 Data sources and workflow

The standard CDR-g workflow was run on all datasets. Brief descriptions of the datasets used are provided below.

#### IFN-beta stimulated human monocytes vs controls

10X genomics. UMI. This is a dataset of two groups of PBMCs. In this experiment, PBMCs were split into a stimulated and control group and the stimulated group was treated with interferon beta.^29^ We utilised a subset of this dataset of CD14+ monocytes.

#### Human muscle dataset

Fluidigm C1. FPKM. Human Skeletal Muscle Myoblasts (HSMM) in either growth or differentiation medium across four timepoints (0, 24, 48, 72 hours).

#### CD8 T cell datasets

Smart-Seq. TPM. CD8 T cells from two datasets^2,30^ of melanoma patients pre and post ICB therapy.

### 2.4 Differential expression analysis

Differential expression analysis was performed using the nonparametric Wilcoxon rank test in SCANPY using log-normalised counts. Cut-offs for log_2_ fold change (LFC) and false discovery rates are described in the text.

### 2.5 Single cell gene set enrichment

Single cell enrichment, which allows the expression profile of CDR-g’s gene sets to be mapped to individual cells, is carried out based on the methodology described in the R singScore package.^26^ Briefly, this method scores gene enrichment by ranking gene expression in individual cells. This method is independent of the gene expression units and scales well to larger datasets. Permutation testing on gene ranks is used to determine whether a gene set is on-or-off in a particular cell. Between conditions, we implement a test of proportions for significance testing, using the Benjamin-Hochberg method for control of false discovery rate.

### 2.6 Functional enrichment

GO-term enrichment on both DEGs and CDR-g’s gene sets utilised GOATOOLS.^31^ This Python package performs functional annotation testing using a hypergeometric test and requires an ontology graph, a list of background genes and a gene-to-GO-term mapping. As input, the GO-slim PANTHER^32^ ontology was used against a background gene set of protein coding genes downloaded from the NCBI’s^33^ gene2go database (dated 2022-01-03).

### 2.7 Code availability

The source code for CDR-g is available at https://github.com/wlchin/pycdr (https://github.com/wlchin/pycdr). The code for dataset analysis can be found in the CDR-g workflow repository (https://github.com/wlchin/CDR_workflows).

## 3 RESULTS

### 3.1 CDR-g is fast, scalable, and outperforms DE-based functional annotation

We first evaluated CDR-g’s performance with synthetic data. For this, we constructed coexpression matrices using a stochastic block model.^22,34^ With this approach, a gene set constituting a gene expression program has a higher “connection” probability amongst its members as compared to other background genes. This allowed us to model gene expression programs that may be shared between conditions but which in the strength of co-regulation between genes, whilst accounting for inherent sparsity of information in single cell data. We planted modules in synthetic co-expression matrices, varying the connectivity strength and the number of conditions. CDR-g efficiently extracted these network modules from 100 different conditions.^22^ Benchmarking CDR-g, we found the algorithm to be efficient, allowing us to investigate 50,000 cells on standard computing hardware (Supplementary figure S1).

Next, we compared CDR-g’s results to genes and pathways discovered by standard DE analysis. We analysed a single cell dataset of interferon-beta stimulated versus unstimulated human peripheral blood monocytes.^29^ We performed DE analysis on this dataset and compared enriched terms from DE genes (absolute log2-fold change (abs log2fc) > 1.0, false discovery rate (fdr) ≤ 0.05) against terms enriched in CDR-g gene sets using the PANTHER GO-SLIM ontology.^32^ With CDR-g, the same enriched term can be represented in multiple CDR-g’s gene sets. Therefore, we compared the DE terms against the union set of all unique terms derived from all CDR-g sets. We found that CDR-g matched 50 of 52 GO terms found by DE (Supplementary figure 2), including the highly relevant terms GO:0034340 (response to type I interferon) and GO:0000165 (MAPK cascade).^35^ Manual inspection of the results revealed that CDR-g did not miss any terms discovered by DE but selected more specific terms (Table 1, Supplementary figure 2). Furthermore, CDR-g detected terms not found by DE that are known to be relevant to interferon biology based on experimental data, including GO:0070534 (K63-ubiquitination)^36,37^ and GO:0007265 (Ras protein signal transduction).^38,39^ These terms were not discovered by DE analysis even when the detection thresholds were relaxed (fdr < 0.1 and abs log2fc cut-off < 0.5).

**Table 1 :**
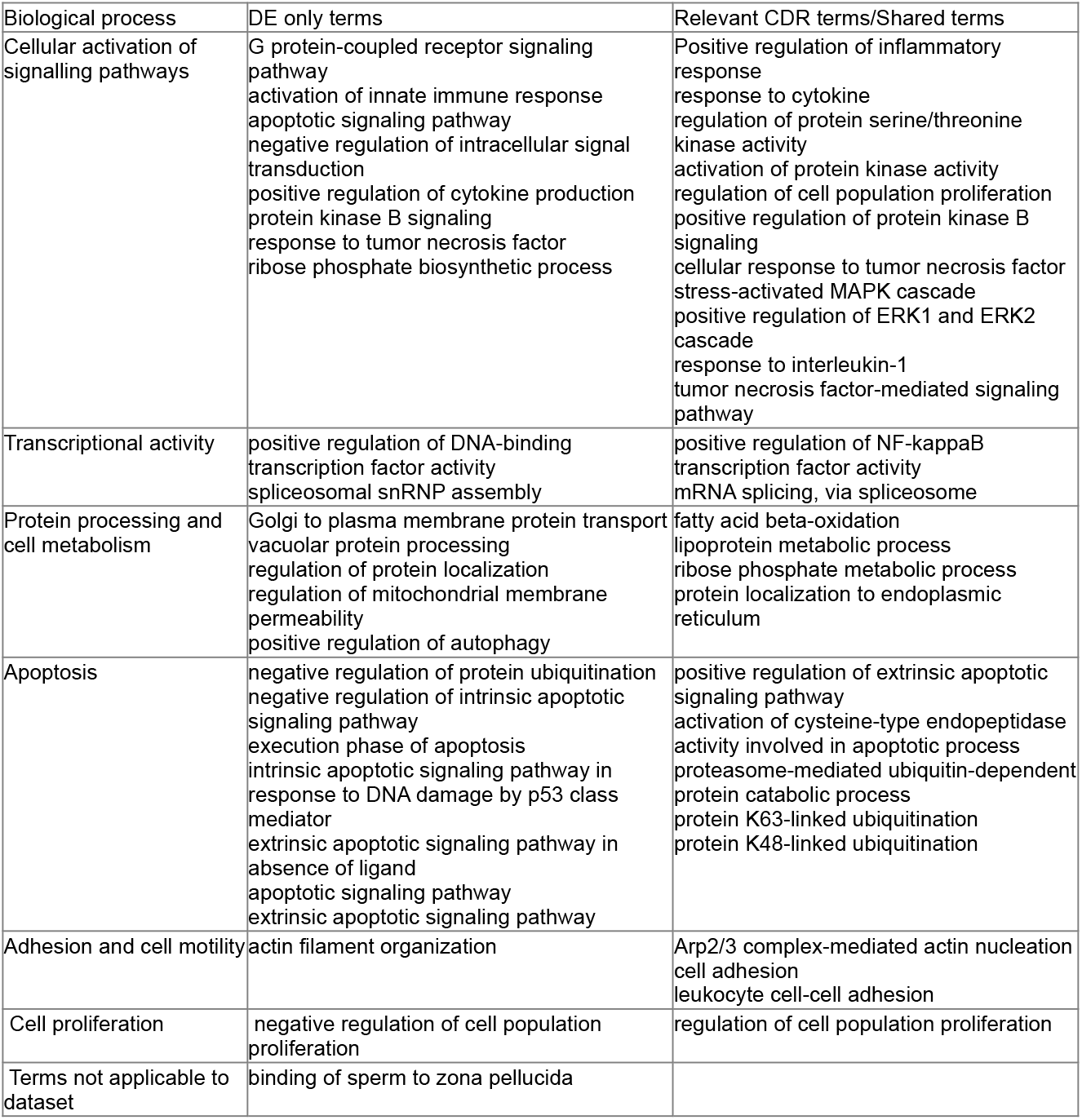
DE only-terms categorised by biological function compared to related CDR-g terms

| Biological process | DE only terms | Relevant CDR terms/Shared terms |
| --- | --- | --- |
| Cellular activation of signalling pathways | G protein-coupled receptor signaling pathway<br>activation of innate immune response<br>apoptotic signaling pathway<br>negative regulation of intracellular signal transduction<br>positive regulation of cytokine production<br>protein kinase B signaling<br>response to tumor necrosis factor<br>ribose phosphate biosynthetic process | Positive regulation of inflammatory response<br>response to cytokine<br>regulation of protein serine/threonine kinase activity<br>activation of protein kinase activity<br>regulation of cell population proliferation<br>positive regulation of protein kinase B signaling<br>cellular response to tumor necrosis factor<br>stress-activated MAPK cascade<br>positive regulation of ERK1 and ERK2 cascade<br>response to interleukin-1<br>tumor necrosis factor-mediated signaling pathway |
| Transcriptional activity | positive regulation of DNA-binding transcription factor activity<br>spliceosomal snRNP assembly | positive regulation of NF-kappaB transcription factor activity<br>mRNA splicing, via spliceosome |
| Protein processing and cell metabolism | Golgi to plasma membrane protein transport<br>vacuolar protein processing<br>regulation of protein localization<br>regulation of mitochondrial membrane permeability<br>positive regulation of autophagy | fatty acid beta-oxidation<br>lipoprotein metabolic process<br>ribose phosphate metabolic process<br>protein localization to endoplasmic reticulum |
| Apoptosis | negative regulation of protein ubiquitination<br>negative regulation of intrinsic apoptotic signaling pathway<br>execution phase of apoptosis<br>intrinsic apoptotic signaling pathway in response to DNA damage by p53 class mediator<br>extrinsic apoptotic signaling pathway in absence of ligand<br>apoptotic signaling pathway<br>extrinsic apoptotic signaling pathway | positive regulation of extrinsic apoptotic signaling pathway<br>activation of cysteine-type endopeptidase activity involved in apoptotic process<br>proteasome-mediated ubiquitin-dependent protein catabolic process<br>protein K63-linked ubiquitination<br>protein K48-linked ubiquitination |
| Adhesion and cell motility | actin filament organization | Arp2/3 complex-mediated actin nucleation<br>cell adhesion<br>leukocyte cell-cell adhesion |
| Cell proliferation | negative regulation of cell population proliferation | regulation of cell population proliferation |
| Terms not applicable to dataset | binding of sperm to zona pellucida |  |

To understand how CDR-g was able to detect novel terms not found by DE analysis, we closely examined all CDR-g gene sets enriched for GO:0070534 (K63-ubiquitination), a term which was not recoverable by DE analysis. In each gene set, CDR-g assigns an z-score rank to each member gene based on permutation importance. Based on this score, top-ranked genes have higher, statistically significant factor loading scores, and therefore are more important members of the gene set. By visualising co-expression matrices alongside CDR-g scores (Figure 2), we confirmed that CDR-g ranked a gene more highly to reflect the degree of differential co-expression between conditions. In all three gene sets enriched for these terms, the five genes driving the enrichment profile (TRAF6, HECTD1, UBE2V2, RNF168, RNF4) were amongst the top 20 genes by Z-score ranking (Figure 2). In these gene sets, the highest ranked genes were also relevant to interferon biology including the transcription factor ATF2,^40, 41^ the interferon-critical co-chaperone CDC37,^42^ and DEFB1 (beta defensin),^43^ further supporting CDR-g’s ability to detect and prioritise key genes within important differential transcriptional programs between interferon-stimulated and unstimulated monocytes.

**Figure 1:**
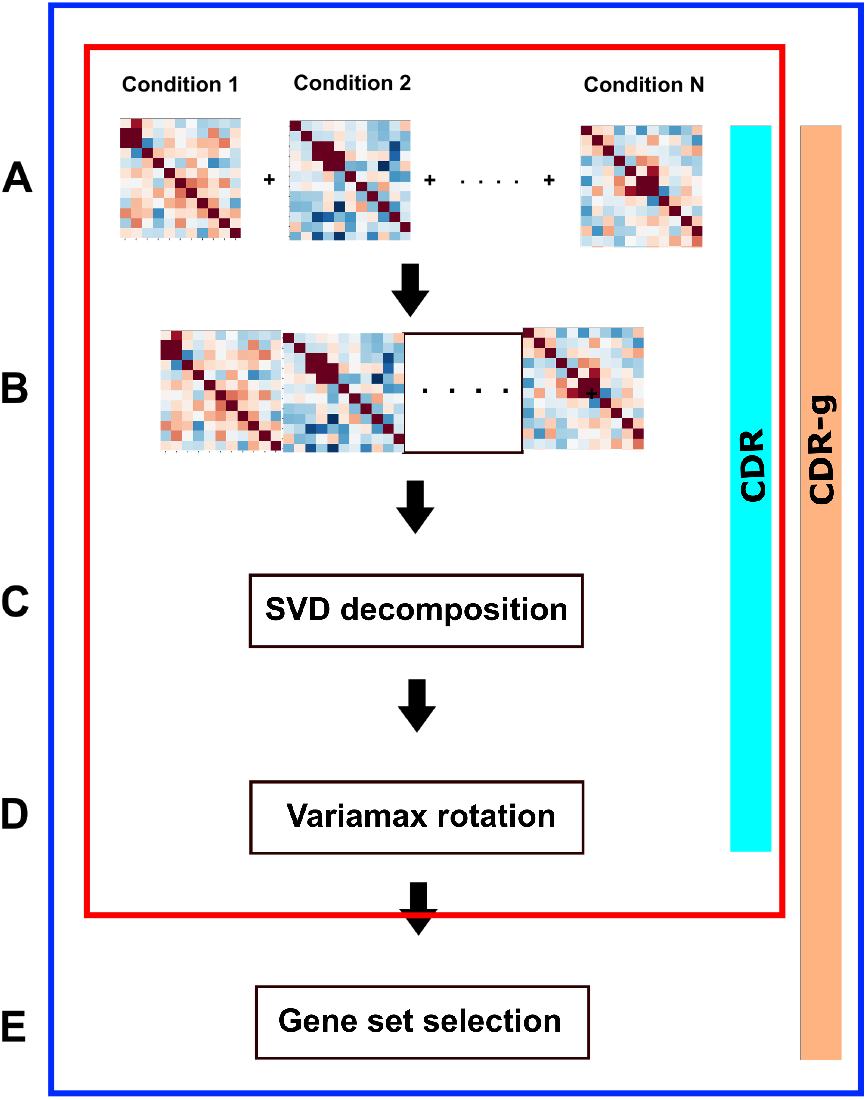
Overview of the CDR-g algorithm. The steps in the CDR-g algorithm involve concatenation of condition-specific gene co-expression matrices (A, B) and the decomposition of their product to produce the resulting factor loading matrix using truncated SVD (C) followed by varimax rotation of the resulting factor loading matrix (D), Gene set selection (E) is performed on each rotated factor loading. Genes which distinguish conditions have high factor loading scores and are selected using a permutation filter. The red box describes the original CDR framework as described in^22^ and the blue box describes CDR-g, the extension to multi-condition single cell data

**Figure 2:**
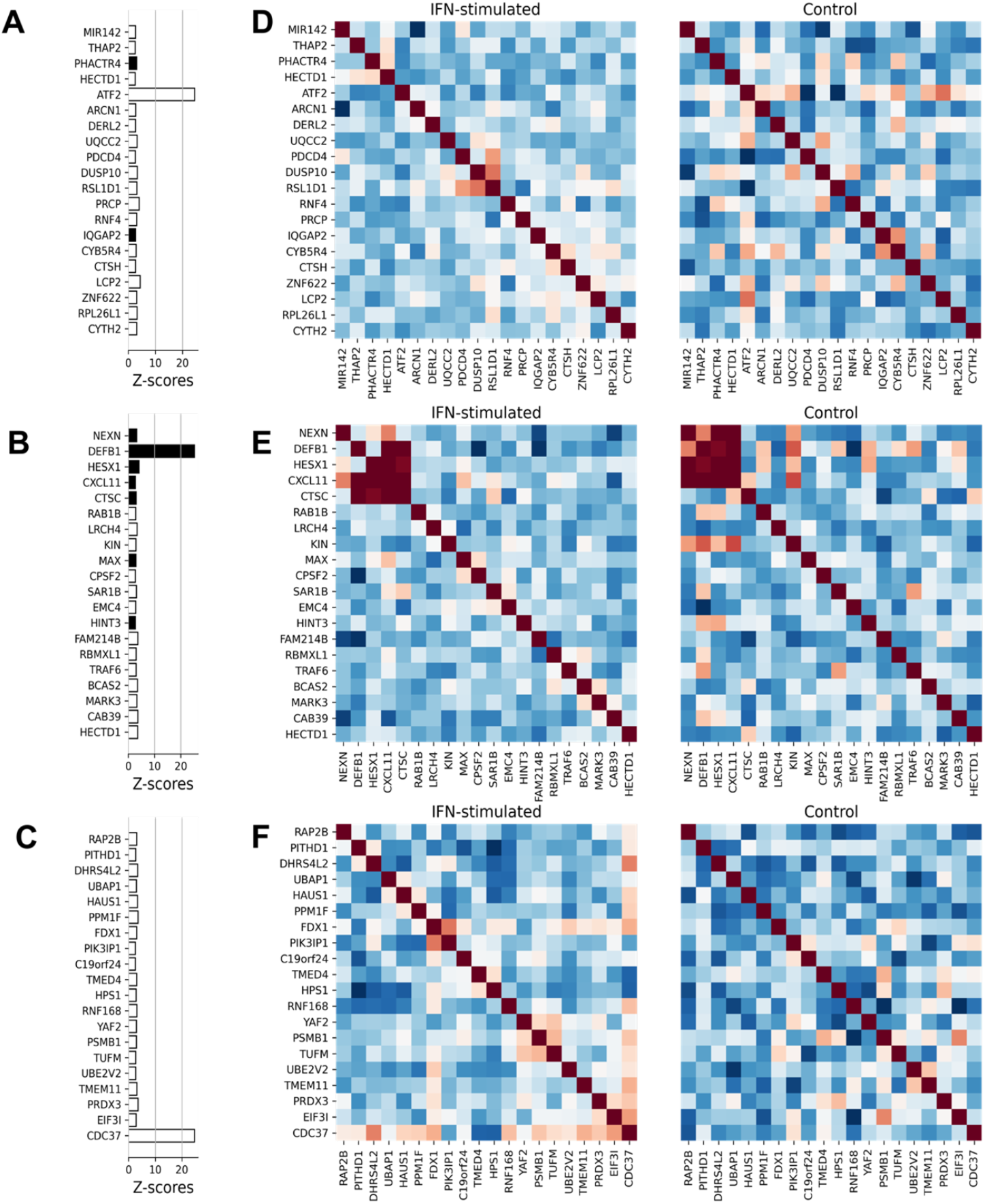
Differentially co-expressed genes in each factor loading account for novel term detection by CDR-g. The rows represent the CDR gene sets enriched for the GO term GO:0070534 (K63 ubiquitination) (A – C): Top 20 genes ranked by Z-score in each CDR gene set. Black bars indicate differentially expressed genes. (D – F): Correlation matrices of top genes for interferonbeta stimulated vs. control monocytes (Red - high correlation; blue - low correlation).

### 3.2 CDR-g identifies transcriptionally distinct subpopulations in time-series single cell data

Next, we tested CDR-g on a single cell RNAseq time-course experiment. The dataset consisted of myoblasts sampled across 4 time points: during the first 24 hours, cells were cultured in a high-mitogen medium to induce myoblast division, followed by three subsequent time points in a low-mitogen medium to induce differentiation.^44^ The original study revealed multiple transcriptional programs superimposed at later time points^44^ and we hypothesized that CDR-g would be able to disentangle these programs. We ran CDR-g using default parameters and uncovered critical pathways by examining terms enriched in CDR-g gene sets. These included GO:0051301 (cell division), GO:0045214 (sarcomere organisation), and GO:0030198 (extracellular matrix organisation) (Figure 3, Table 2). The proportion of cells active for these gene sets at each time point corresponded to the switch in incubation medium (Supplementary figure 4). CDR-g also yielded additional genes, critical to these expression programs without requiring further bioinformatic analysis (Supplementary figure 4). CDR-g identified TGF*β*I,^45^ a gene influencing differentiation of myotubes in cultured human myocytes, LMOD3,^46^ an AKT and ERK-dependent myogenic regulator and DNA replication licensing factors^47^ such MCM4 and MCM6 amongst the top ranked genes from these expression programs.

**Figure 3:**
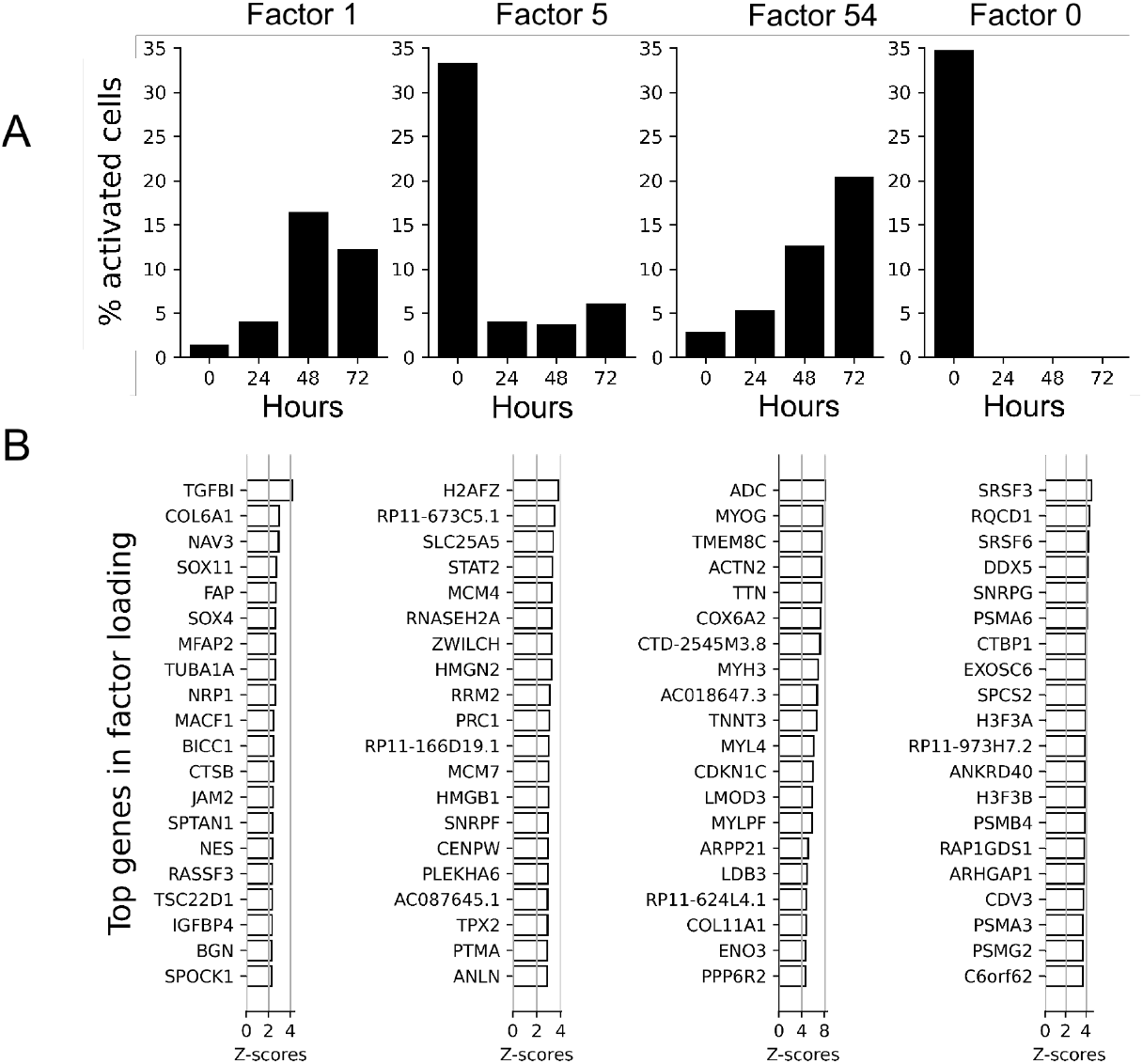
Enrichment profile for key CDR-g gene sets across experimental timepoints in muscle dataset. (A) Proportion of cells showing gene set activation across time point, as measured using single cell enrichment. (B) Key genes from each gene set, as ranked using z-score. Terms enriched in each gene set are provided in Table 2.

**Table 2 :** Enriched terms in each CDR-g gene set described in the muscle data analysis in the main text.

| Factor loading | Gene sets |
| --- | --- |
| Factor loading 8 | extracellular matrix organization<br>skeletal muscle tissue development<br>wound healing<br>cellular zinc ion homeostasis<br>cellular response to transforming growth factor beta stimulus',<br>cell adhesion<br>skeletal system development<br>response to metal ion<br>cell-matrix adhesion<br>collagen fibril organization |
| Factor loading 7 | cell division<br>kinetochore assembly<br>DNA repair<br>chromosome segregation<br>mitotic cytokinesis<br>mitotic sister chromatid segregation<br>G2/M transition of mitotic cell cycle |
| Factor loading 1 | extracellular matrix organization<br>cell adhesion<br>wound healing |
| Factor loading 5 | cell division<br>chromosome segregation<br>DNA replication<br>DNA repair<br>cellular response to DNA damage stimulus<br>mitotic cytokinesis<br>interstrand cross-link repair<br>kinetochore assembly<br>mitotic sister chromatid segregation<br>mitotic cell cycle phase transition<br>DNA replication checkpoint<br>mitotic cell cycle |
| Factor loading 54 | sarcomere organization<br>muscle contraction<br>skeletal muscle contraction<br>actin filament organization<br>cardiac muscle cell development<br>cardiac muscle contraction |
| Factor loading 0 | negative regulation of G2/M transition of mitotic cell cycle<br>NIK/NF-kappaB signaling<br>regulation of mitotic cell cycle phase transition',<br>mRNA processing<br>protein folding<br>regulation of alternative mRNA splicing, via spliceosome<br>RNA export from nucleus<br>mRNA export from nucleus<br>tumor necrosis factor-mediated signaling pathway<br>proteasome-mediated ubiquitin-dependent protein catabolic process<br>SCF-dependent proteasomal ubiquitin-dependent protein catabolic process<br>positive regulation of canonical Wnt signaling pathway<br>mRNA 3'-end processing<br>negative regulation of canonical Wnt signaling pathway<br>post-translational protein modification<br>protein deubiquitination<br>RNA splicing, via transesterification reactions<br>T cell receptor signaling pathway<br>translational initiation<br>mitochondrial translation<br>protein polyubiquitination<br>regulation of mRNA splicing, via spliceosome<br>translation<br>mRNA cis splicing, via spliceosome<br>regulation of translation<br>MAPK cascade<br>protein stabilization |

Terms enriched in CDR-g gene lists (Figure 3, Table 2) included SCF-mediated ubiquitination 44 and Wnt signalling,^48^ pathways related to myogenesis which we could not elicit from this dataset using pairwise DE. Interestingly, we found that these biologically important terms mapped not only to the same time point (time point zero), but to a single gene set (Factor loading 0, Figure 3A, Table 2), hinting that this gene set contained key early myogenic regulators. We hence examined the ranked gene list from this gene set. PSMA3, PSMB4 and PSMA6, are protea-some genes crucial to muscle development.^49^ DDX5 is a cofactor of MyoD^50^ and CTBP1 is a transcriptional co-repressor that binds to the myogenin promoter. H3.3 is incorporated in the regulatory regions and promoters of myogenic genes prior to differentiation and its recruitment increases during differentiation.^51^

From our examination of terms enriched in the first two time points (0 and 24 hours) by CDR-g analysis, we also identified a gene expression program (Figure 4) which displayed maximal activation at time point 0. Interestingly, this gene expression program was enriched for terms including extracellular matrix organisation (GO:0030198) and cell-matrix adhesion (GO:0007160) (table 2). We expected time-point zero to contain only rapidly dividing myoblasts grown incubated in high-mitogen medium, yet these terms represent pathways found in mature, differentiating myocytes. We postulated that these terms could be accounted for by subpopulations of cells displaying divergent transcriptional programs at time point zero. To test this, we examined the factor loading accounting for this divergent program using single cell enrichment and found that activation of this gene expression program present in non-dividing cells (Figure 3). Amongst the top genes in this factor loading was the homeobox protein PRRX1,^52^ TGFBI,^53^ FBLN5,^54^ and LTBP1,^55^ pointing to a TGF beta-centric mesenchymal program in these cells. Previously, it was reported that a population of cells displaying mesenchymal-like properties was seen at time point zero and almost all cells were thought to display proliferating properties based on CDK1 staining.^44^ CDR-g resolved timepoint zero myocytes in this dataset into two distinct populations based on mitotic activity and differential pathway activation, in addition to providing enhanced resolution of potential transcriptional programs that were normally not detected using conventional DE analysis.

**Figure 4:**
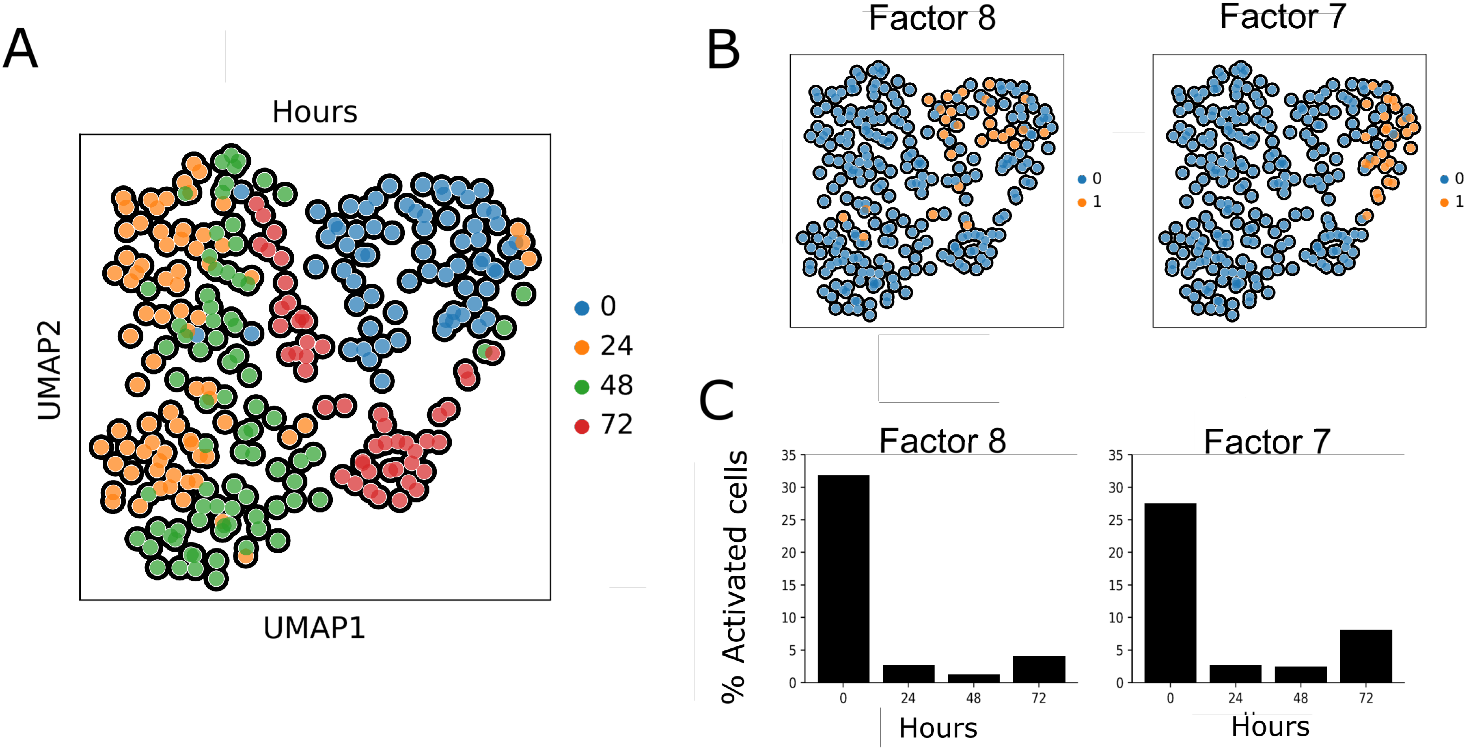
CDR-g resolves cells at timepoint zero into two populations based on transcriptional activity. (A) UMAP of myocytes colored by timepoint. (B) Percentage of cells showing activation of gene sets at each time point, with their corresponding UMAP diagrams in (C). Factor loading 8 is a gene set enriched of TGF-beta pathway terms . Factor loading 7 is a gene set enriched for cell division terms. Terms enriched in each gene set are provided in Table 2.

**Figure 5:**
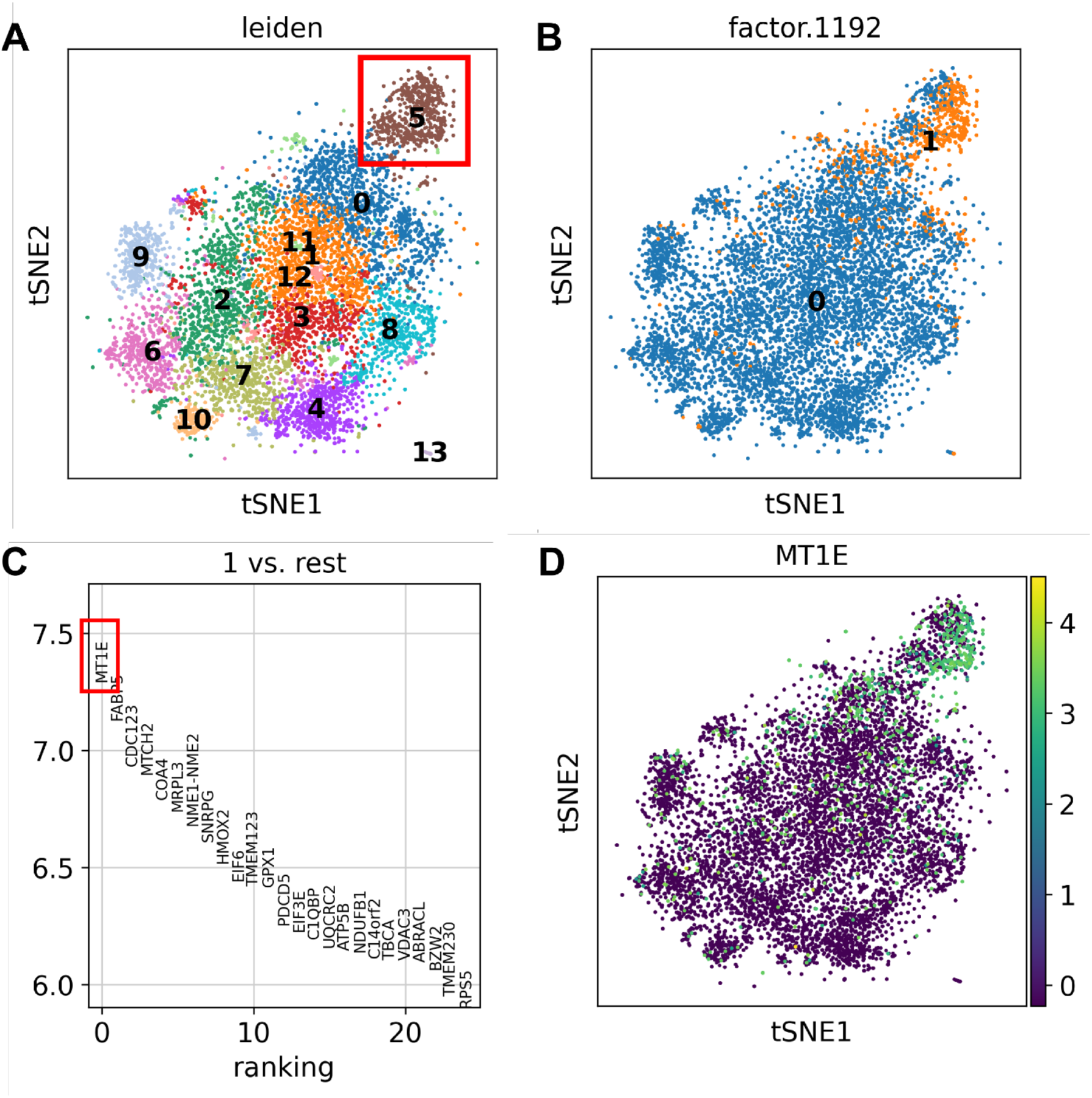
CDR identifies an MT1E-enriched subcluster of dysfunctional T cells. (A) tSNE of combined dataset with actively dividing CD8+ T cells highlighted (red box, cluster 5). (B) shows cells displaying activation of CDR hypoxia gene set (factor.1192) in orange. (C) Conserved marker analysis on cluster 5, partitioning by CDR gene set activation. (D) MT1E gene expression in combined dataset.

### 3.3 CDR-g identfies transcriptionally distinct subpopulations from combined single-cell datasets

Integrating single cell data sets from multiple studies is becoming increasingly important. We therefore tested CDR-g on a combined data set from two distinct melanoma studies^2,30^ composed of immune cells from patients at various stages of checkpoint blockade therapy. Given the importance of CD8+ T cells in the immune checkpoint blockade (ICB) response,^56^ we focused on this population of cells in our analysis, using cells labelled as CD8+ T cells from each of their original dataset. In total, this combined dataset was composed of 8109 CD8+ T cells across 48 patients, sampled either before or after treatment with ICB therapy. Graph-based clustering at default resolution revealed 14 single cell clusters. (Figure 4). Rather than individually analyzing these clusters, we hypothesized that CDR-g could summarise key sources of differential variation in this dataset and hence offer a summary of CD8+ T cell transcriptional programs in melanoma.

We combined expression matrices from the two datasets and supplied only clinical phenotype information in the two datasets to CDR-g. We examined the union set of enriched terms produced by CDR-g’s gene sets. We discovered terms belonging to five categories (cell proliferation and division (GO:0051301), lymphocyte activation and cytolytic activity (GO:0042110), interferon signalling and cytokine pathways (GO:0060337), metabolic processes (GO:0051248) and protein processing pathways (GO:0061077)). Representative gene expression programs are shown in supplementary figure 3. These processes form distinct superimposed activation patterns across the combined dataset, crossing the cluster boundaries defined by conventional graph-based clustering.

We observed that the cluster composed of proliferating T cells (Figure 4, Supplemental Figure S3) could be further subdivided based on a CDR-g gene set activation profile (cellular response to hypoxia, GO:0071456). Prior analysis,^56^ which used a hierarchical clustering strategy based on marker expression to distinguish T cell subclusters, had labelled this population as single population of exhausted but dividing T cells. Therefore, we analysed conserved markers in this cluster in more detail to further delineate these subpopulations. Partitioning this subcluster based on CDR-g gene set activation showed a smaller subpopulation of T cells expressing MT1E as a conserved marker. MT1E is a unique experimentally validated marker for T-cell dysfunction,^57^ uncoupled from typical markers of T cell dysfunction such as TIGIT and PCDC1. This information could not be recovered through a graph-based clustering, even at high resolution (Supplemental figure S4). Our analysis illustrates the value of CDR-g in identifying discrete yet biologically relevant differences in cell states and summarising pathway level shared between datasets, thus improving, and simplifying data interpretation from complex multi-variable experiments.

## 4 Discussion

The number of complex single cell datasets continues to grow year after year, as do the number of methods used to analyse such data. The rapid pace of development of analytical strategies for single cell data means that practitioners are confronted with a plethora of methods designed to derive biological insight from complex multi-condition experimental designs. With CDR-g, the ability to investigate different aspects of transcriptional variation in a single, unified computational framework is valuable to researchers working with single cell data. We have shown that CDR-g can be applied to a variety of different research contexts and that it scales well.

More recently, new methods have been designed for complex multi-condition data including Milo^58^ which examines differential abundance, GDMF,^30^ which examines shared variation and Augur,^59^ which allows cell-type prioritisation of multi-condition perturbations. CDR-g distinguishes itself from these methods by directly considering the coexpression structure across conditions of interest, providing a robust way to characterize this important aspect of transcriptional variation in a single cell dataset.

Finally, we address some practical aspects of running a CDR-g analysis. Because CDR-g applies SVD and varimax rotation as part of its algorithm, CDR-g will prioritise the largest variances in differential co-expression across conditions. Technical variation and batch effects may be captured by CDR-g if these elements contribute to variation between conditions, so data pre-processing and data normalisation is required. Additionally, although we have shown that sub-populations within conditions can be detected using a CDR-g analysis, rare cell populations may escape CDR-g’s detection threshold because their signal is subsumed during condition-specific co-expression matrix construction. If finer resolution is desired, constructing co-expression matrices on smaller cell subsets (i.e. by sub-partitioning conditions of interest using finer cluster parameters) can enhance biological pathway detection by CDR-g.

In summary, we present an efficient, novel approach for identifying gene expression programs across multiple conditions designed for large and complex single cell datasets. We anticipate that its broad applicability to different study designs combined with its computational properties will be of great benefit to the community.

## 5 Authors’ contributions

L.P and M.S developed the mathematical theory and formulated the CDR algorithm. L.P and W.L.C wrote the implementation of the algorithm. T.L and W.L.C extended the algorithm to multicondition single cell data. W.L.C wrote the manuscript and the analysed the human datasets. W.J.L, M.S, L.P and T.L reviewed the manuscript.

## 6 Data Availability

The datasets used for analysis have been described in the Materials and Methods section.

